# Nucleosome assembly and disassembly pathways

**DOI:** 10.1101/2020.04.01.020263

**Authors:** A. Hatakeyama, R. Retureau, M. Pasi, B. Hartmann, C. Nogues, M. Buckle

## Abstract

Nucleosome assembly and disassembly play a central role in the regulation of gene expression. Here we use PhAST (Photochemical Analysis of Structural Transitions) to monitor at the base pair level, structural alterations induced all along DNA upon histone binding or release. By offering the first consistent, detailed comparison of nucleosome assembly and disassembly *in vitro*, we are able to reveal similarities and differences between the two processes. We identify multiple intermediate states characterised by specific PhAST signatures; revealing a complexity that goes beyond the known sequential events involving (H3-H4)_2_ tetramer and H2A-H2B heterodimers. Such signatures localise and quantify the extent of the asymmetry of DNA/histone interactions with respect to the nucleosome dyad. This asymmetry is therefore defined by the localisation and amplitude of the signals. The localisation of the signal is consistent between assembly and disassembly and dictated by the DNA sequence. However, the amplitude component of this asymmetry not only evolves during the assembly and disassembly but does so differently between the two processes.

Understanding the regulation of gene expression requires a complete knowledge of nucleosome dynamics. Our unexpected observation of differences between assembly and disassembly opens up new avenues to define the role of the DNA sequence in these processes. Overall, we provide new insights into how the intrinsic properties of DNA are integrated into a holistic mechanism that controls chromatin structure.

**Statement of Significance:** This manuscript addresses the question of nucleosome dissociation compares with association. We used PhAST which is a non-intrusive photochemical technique to follow nucleosome dynamics at base pair resolution. We observed structural asymmetry during nucleosome turnover. We also showed for the first time that the process of nucleosome dissociation is not a reversal of association. This asymmetry favours intermediate states involved in chromatin organisation suggesting novel models for the role of nucleosome turnover in the epigenetic regulation of gene expression.

## Introduction

The fundamental repeating unit of chromatin is the nucleosome; 145-147 base pairs (bp) of DNA wrapped around an octamer of histone proteins (two H2A/H2B heterodimers and one (H3-H4)_2_ tetramer) that efficiently compacts genomes into cell nuclei and regulates many DNA functions. The spatial and temporal distribution of nucleosomes as a result of nucleosome assembly and disassembly is involved in all DNA transactions. *In vivo*, a plethora of interplaying factors such as chaperones, remodelling complexes, histone variants, epigenetic modifications and intrinsic, sequence dependent, DNA properties orchestrate the dynamics of nucleosome positioning. A detailed knowledge of the biophysical basis underlying the structural pathways involved in nucleosome assembly and disassembly is a prerequisite to understanding such an important and fundamental cellular event as nucleosome biogenesis and turnover.

In recent years a number of studies have been conducted on the mechanism of nucleosome disassembly. Studies were carried out *in vitro* by recording the response of preformed nucleosomes to a gradual increase in ionic strength [1, 2] or, less frequently, to an external mechanical force applied to the nucleosome DNA [3, 4] [5]. The examination of spontaneous disassembly focused on unwrapping of DNA in the peripheral regions (so-called “DNA breathing”) [6–10]. Fluorescence-based techniques were used to dissect the stepwise disassembly of nucleosomes, each experiment providing data about specific histone-histone, DNA-histone or, eventually, DNA-DNA proximities. For example, the use of three or four pairs of fluorophores gave insights into the behaviour of the DNA’s extremities [1] or interior DNA regions [11] with respect to H4 and H2B, as well as providing information about histone-histone proximities [12].

In theory, two global pathways for nucleosome disassembly may be envisaged, either dissociation of the octamer as a single entity from the DNA or sequential release of histones or groups of histones; the second hypothesis is now generally accepted. A series of studies of salt-induced dissociation based on FRET (Forster Resonance Energy Transfer) approaches [1], 2, 11, 12, [11–14] [2) or TR-SAXS (Time-Resolved Small Angle X-ray Scattering) {Chen, 2014 #3253, 15, 16] [17] provided coherent arguments in favour of a pathway with two major successive phases: a first step leading to the release of H2A-H2B dimers and a second, distinct step corresponding to (H3-H4)_2_-DNA dissociation. Such a global two-step scheme is likely to be general since it was observed in nucleosomes studied under identical conditions but containing different DNA sequences, *i.e*. 601- and 5S-nucleosomes [13, 16] or 601-, 5S- and MMTV-nucleosomes [14, 18].

Although technically not straightforward, the existence of very early states of disassembly, before the removal of H2A-H2B, was also examined. With the 5S-nucleosome, the two H2A-H2B dimers were observed to dissociate from the (H3-H4)_2_-DNA complex in a single transition without observable stable intermediates [1] [16]. FRET experiments with 601-nucleosomes proposed that disruption or weakening of the interface between H2A–H2B dimers and the (H3–H4)_2_ tetramer (the so-called “butterfly” state), helped to rupture the DNA / H2A–H2B interfaces [11] [2, 14]. According to models inferred from experiments using SAXS [16], FRET [4] and single molecule unwrapping associated with FRET [4], the release of the two H2A–H2B dimers in 601-nucleosomes is asymmetric, starting from one unwrapped DNA end (the so-called “J”-shaped state). A recent cryo-EM study captured structures related to first events of spontaneous disassembly of 601-nucleosomes [10], which are hard to observe in solution because nucleosome open states are marginally populated [6] [7]. The cryo-EM structures showed again an asymmetric loss of contacts between H2A-H2B and DNA arising from the spontaneous breathing of one extremity of the DNA fragment; this first nucleosome opening gradually propagates in association with subtle histone rearrangements; the intermediate states, in which one H2A–H2B dimer is no longer visible, resemble those hexasomes (DNA bound to the (H3–H4)_2_ tetramer and one H2A–H2B dimer) obtained from SAXS [19] or X-ray [20] data. In all the studies presented above, as well as in a more recent study using SAXS [21], the asymmetric opening was considered symptomatic of a DNA sequence effect since the strict symmetry of the histone structured domains with respect to the dyad axis [22] [23] cannot account for such phenomena. On the basis of salt titrations [22] and single molecule experiments [3], it was proposed that the 601 sequence is constituted by “strong” left and “weak” right halves [17]; strong and weak sides may relate to differences in DNA intrinsic flexibility [24].

A general DNA sequence effect on disassembly was further attested by the fact that 601-nucleosomes better resist chaotropic destabilisation than nucleosomes formed with other sequences [13] [21]. In contrast, nucleosome stability, characterised by the ionic strength at the midpoint of the disassembly transition, is not affected by the histone origin [14] probably because of the very high degree of conservation of histone sequence and folding [25].

The large number of studies presented above provided information exclusively on nucleosome disassembly; states preceding H2A–H2B dimer release are better characterised than later events leading to the complete dissociation of the complex. By comparison, assembly remained almost completely ignored apart from initial studies during the 1980’s [26] [27], a situation that prevented any global approach to understanding both nucleosome association and dissociation. In view of this, we embarked on studies to capture nucleosome intermediate states during NaCl-induced assembly and disassembly (Schema 1) of the 601-nucleosome and to compare the two pathways. We used the PhAST (Photochemical Analysis of Structural Transitions) technique developed in our group [28].

PhAST possesses the useful property of being able to follow both processes from the DNA point of view and to provide structural information at base pair resolution, independent of conditions. This non-invasive technique can thus be applied to freely diffusing macromolecules in solution, without introducing structural artefacts caused for instance by bulky hydrophobic fluorophores [12]. It is based on the measurement of the probability to form, on the same DNA strand, UV photo-induced cyclobutane dimers between adjacent pyrimidines (YpY dimer, linked by C5-C5 and C6-C6 bonds) [28–30]. In addition to the quantum yields specific to each type of step, the probability of forming YpY dimers depends on the local YpY structure; more precisely on the inter base-pair parameters of roll and twist that are coupled in B-DNA in solution [31]. Thus, low twist and positive roll shorten the YpY C5-C5 and C6-C6 distances and thus favour dimer formation whereas large twist and negative roll have the inverse effect [28]. The YpY dimer probabilities along a DNA sequence therefore reflect its local average structure. Comparison between probabilities collected on free and bound DNA simultaneously reveals DNA structural changes induced by the presence of proteins.

PhAST proved to be remarkably efficient in following structural changes in DNA as 601- and derived 601-nucleosomes were formed under decreasing ionic strength conditions, *i.e*. during nucleosome assembly [28]. We demonstrated first, the intuitive idea that histone binding induces noticeable structural changes in the local parameters of roll and twist all along the 601 sequence. Then we observed that nucleosome formation starts with the binding of (H3/H4)_2_ and ends with the recruitment of H2A-H2B dimers, in agreement with earlier studies [26] [27]. An important original contribution of PhAST was to detect and describe additional steps occurring during nucleosome formation. The structural organisation of nucleosome intermediate states reflected the existence of marked DNA sequence effects that could be unambiguously assigned to specific DNA regions. For example, (H3/H4)_2_ interacts more robustly with the 5’ side of the 70bp central segment of the 601 sequence than with its 3’ side counterpart, a dissymmetry that is not present when the 5’ side is mutated at key points. These results were explained by the experimental asymmetries found in the dynamic properties of the free sequence [24]. That such subtle events could be captured by PhAST encouraged us to further describe how the 601-nucleosome disassembles allowing us to compare assembly and disassembly processes.

In this manuscript we first examined the potential effect of high salt concentrations to induce structural perturbations in free DNA, [32] by examining data collected on the free 601 sequence at various ionic strengths and comparing the YpY reactivities. We then catalogued differences in YpY reactivities between free and bound DNA thus obtaining a description of DNA structural changes during increased or decreased ionic strength. To interpret the structural changes in the local DNA configuration in terms of strengthening or weakening of DNA/histone interactions (ultimately, histone binding or release), we used a very fine mapping of contacts between DNA and both structured and unstructured histone regions obtained from exhaustive molecular simulations in explicit solvent [33]. A parallel between DNA structural changes and DNA/histone interactions led to the identification and characterisation of a series of predominant nucleosome intermediate states. Finally, we discussed how the structural organisation of the intermediates that constitute the assembly and disassembly pathways relate to the properties of free DNA.

## Materials and Methods

### Nucleosome reconstitution

DNA fragments containing 601 sequence were prepared as described previously [28]. Nucleosomes were reconstituted with a salt dilution method according to manufacturer’s instructions (New England BioLabs) with a slight modification. Human recombinant histone H2A/H2B dimer (1.5 μg, 54 pmol) and histone (H3/H4)_2_ tetramer (1.5 μg, 27 pmol) (New England BioLabs) were mixed with the linearised 601 DNA fragments (6 μg, 16 pmol) in 11 μl of 2M salt buffer (18 mM Tris-HCl, 2 M NaCl, 0.9 mM DTT, 0.9 mM EDTA). The mixture was incubated at room temperature (RT) for 30 min before the salt concentration was lowered to 0.1 M by adding dilution buffer (10 mM Tris-Cl, pH 7.5, 1 mM EDTA, 0.05% NP-40, 5 mM 2-mercaptoethanol, 0.1 mM PMSF) five times every 20 min (from 1.5 M to 1.0 M, 0.5 M, 0.25M, and 0.1M NaCl).

### Dissociation process

Preformed nucleosome complexes were dissociated using two different regimes of increasing salt concentration; either nucleosomes or free DNA were transferred stepwise from 0.1 M NaCl to 0.5 M, 1.0 M and 1.5M. After 15 minutes incubation at each of the salt concentrations, samples were photoirradiated with the laser as described below. Alternatively, nucleosomes or free DNA at 0.1 M NaCl were transferred directly by one step to either 0.5 M, 1.0 M or 1.5 M NaCl and irradiated with the laser. As both regimes led to almost identical results (supplementary data Figure S1), the data analysed here are from the two types of experiments considered together.

#### One step salt increase

Twenty microlitre (20 μl) of preformed nucleosome complexes prepared as described above (30 ng/μl DNA) was mixed with 20 μl of high salt buffer (10 mM Tris-Cl, pH 7.5, 1 mM EDTA, 0.05% NP-40, 5 mM 2-mercaptoethanol, 0.1 mM PMSF at one of the following NaCl concentrations; 1M, 2M, and 3M). The final NaCl concentration was 0.5 M, 1 M, or 1.5 M. Final sample volume was 40 μl (DNA concentration was 15 ng/μl) for all conditions. The mixture was incubated for 20 min at RT before irradiation with the laser.

#### Stepwise salt increase

One hundred microlitres (100 μl) of preformed nucleosome complexes in 0.1 M NaCl buffer prepared as described above (30 ng/μl DNA) was mixed with 11 μl of 4M salt buffer (10 mM Tris-Cl, pH 7.5, 4M NaCl, 1 mM EDTA, 0.05% NP-40, 5 mM 2-mercaptoethanol, 0.1 mM PMSF) to increase the salt concentration to 0.5 M and the mixture was incubated for 20 min. 4M NaCl salt buffer was added in two more steps to increase the NaCl concentration from 0.5M to 1.5M. The mixture was incubated for 20 min after every addition. At each salt concentration, 30 μl of the mixture was taken for PhAST analysis. DNA concentration was adjusted to 15 ng/μl by adding buffer with NaCl before irradiated with the laser.

### Association process

A series of PhAST experiments were previously carried out to study the nucleosome assembly as described [28]. As for the disassociation described above, the experiments were carried out in two ways; either preformed nucleosomes were submitted to a progressive decrease of the ionic strength, or a mixture of free DNA and histones were transferred directly by a single step to different NaCl concentrations. Both approaches gave similar results, and the published data, reused here, are from the two types of experiments considered together. We also ensured that the assembly process followed the same pathway as described in the original paper so that our use of previous data is justified.

### PhAST analysis

Samples of DNA alone or of reconstituted nucleosomes were irradiated with the UV Laser, followed by primer extension of fluorescent-end-labelled oligonucleotides and separation of the ensuing fragments using capillary electrophoresis (CE, for details on the experimental procedure see [28]). The resulting electrophoretograms (see Figure 1 of our previous publication [28] for an example) were analysed to determine the size (in base pairs) and relative abundance of the fragments present in each sample, using the following procedure. First the electrophoretogram was calibrated by converting migration times to fragment sizes (in units of bases) through piecewise linear fits to the internal size standard (600LIZ) which was run together with each sample. The initial part of each electrophoretogram (up to sizes corresponding to 20 nucleotides) consistently displayed extremely noisy behaviour and was systematically discarded from further analyses. Note that since the primers used for primer extension were larger than 20 b (see above), the retained portion of electrophoretograms also contained the unelongated primers. To facilitate comparison among independent CE runs, electrophoretograms were then normalised using their integral. Peaks were identified based on the analysis of the numerical first derivative of the electrophoretograms, and their maximum height was taken as an estimation of the relative abundance of each fragment. Knowing that a fragment of length *x* indicated the formation of a pyrimidine dimer between bases *x* + 1 and *x* + 2, to obtain the relative propensity of forming a pyrimidine dimer at each position along our DNA, we needed to assign to each peak an integer fragment size in units of bases. The starting point for this assignment was the calibrated migration time corresponding to the peak maximum, which was, by the very nature of the calibration process, a fractional quantity in units of bases. Instead of simply taking the closest integer size by rounding, which often leads to artefacts such as assigning two peaks to the same fragment size, we developed an optimization procedure that minimised artefacts by allowing small corrections (on average of about 0.24 bases in either direction), rigorously without changing the order of peaks. In particular, we took advantage of the fact that the size of a fragment implies its sequence, and made the reasonable assumption that the most likely cause for the polymerase to stop is the presence of a photo-induced pyrimidine dimer. Our procedure maximizes, within the aforementioned constraints, the likelihood of the fragment length assignment given its sequence. The resulting sets of peak intensity as a function of DNA sequence for *n* independent replicates at the same salt concentration were averaged (*n* = 3 for the association data as in our previous paper [28], *n* = 6 for the dissociation data collected here), and the standard error was calculated.

To quantify changes in the likelihood of pyrimidine dimer formation between free and bound DNA at a given salt concentration, we calculated the ratio between the peak intensity in bound DNA over free DNA for each pyrimidine dinucleotide on both strands. These ratios indicate how much more or less likely it is for a given pyrimidine dimer to form in the presence of histones. As described previously [28], the comparison is best presented using the log_2_ of these intensity ratios (log_2_IR, see e.g. Figure 3). The standard error σ for the log_2_IR values was estimated using the following formula:

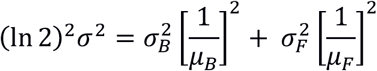

which propagates the standard errors calculated for each condition (*σ_B_* and *σ_F_* for bound and free DNA, respectively) using a first order expansion around their means (*μ_B_* and *μ_F_*).

At a very few positions, PhAST signals corresponded to weak reactivities that were so variable across the experiments that log_2_(*IR*) values were lower than the associated standard error; these suspicious points were discarded from the analysis illustrated in Figures 5 and 6.

### Nucleosome models, simulations and DNA/histone interface analysis

The construction of the nucleosome models, the set-up of simulations and the DNA/histone interface analysis were detailed in an original article recently published [33]. We present a brief summary to place the analyses presented here in their initial context.

The four studied nucleosome models contain *Xenopus laevis* histones and the 601 sequence. All models included the histone tails that were partially truncated to only retain those regions not digested by trypsin and clostripain [34] [35]. These nucleosomes were simulated by molecular dynamics in explicit solvent (water molecules and 150 mM NaCl) using the CHARMM36 force field [36] for a total duration of 1μs.

The interface between DNA and histones was analysed by VLDM (Voronoi Laguerre Delaunay for Macromolecules), a software used in particular to depict the DNA/protein interfaces without resorting to any empirical or adjusted parameters [33] [37]. The interface between two structural elements is a polygonal surface, quantified by its area and by its occurrence. In this paper, the contacts occurring less than 20% of the simulation time were not considered.

Note that the DNA sequence is expressed in terms of Super Helical Location (SHL) that is, the number of helical turns separating a given base pair from the central base pair, SHL0; we assume that, on average, one turn corresponds to 10 bp.

## Results

PhAST generates YpY dimers in DNA using laser photo-radiation; the dimer detection technique produces peaks representing the probabilities of dimer formation; the quantification consists of measuring the peak amplitudes which we will call intensities (*I*) (for calculation of these see Materials and Methods). The intensities (*I*) reflect the DNA local structure, as reported in the Introduction. In the two possible end situations, intensities (*I*) correspond to i) free DNA or ii) DNA fully engaged in a nucleoprotein complex. Intermediate values characterise nucleosome intermediate states. By comparing data collected throughout nucleosome assembly and disassembly with free DNA, PhAST therefore reveals the structural effects of protein binding on DNA.

Details of the PhAST experiments are given in Materials and Methods. Broadly speaking, they follow the Schema 1, using either decreasing or increasing ionic strengths to study assembly or disassembly, respectively. Dissociation was carried out either by stepwise salt increase or one step alst increase (see Materials and Methods). No difference in result was observed between the two techniques (Supplementary Figure S1). From Schema 1 note that the intensities (*I*) corresponding to assembly were previously published [28] and reused here for comparison. In all cases, free DNA was in parallel photoirradiated in the absence of histones.

**Schema 1:**
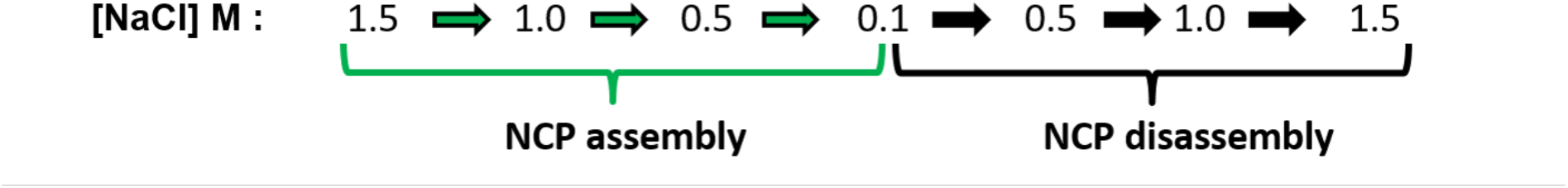
Outline of the PhAST experiments monitoring successive DNA structural changes induced during nucleosome assembly and disassembly by using stepwise decrease and increase of NaCl concentration.

Before presenting our analysis of differences in PhAST signals associated with variations in salt concentration and their interpretation in terms of changes in DNA/histone interactions, we focus on the free DNA to ensure that ionic strength variations in the range used here do not perturb its structure.

### PhAST profiles of free DNA at various NaCl concentrations

Our studies of nucleosome assembly and disassembly used salt concentrations that range from 0.1 to 1.5 M NaCl. Variations within this range modify the DNA melting temperature [38] without strongly perturbing the gyration radius or persistence length of B-form of DNA [39–41]. However, to our knowledge, there is no report on possible effects of such moderate ionic strength variations on the local structure of the B-DNA double helix at room temperature. We decided to examine the effects of salt concentration on the PhAST signals of DNA alone to obviate any potential bias on interpretation of the data. We therefore compared the PhAST signals collected on the 601 sequence at 0.5, 1.0 or 1.5 M NaCl to those obtained at 0.1 M NaCl (Figure 1). If normalised intensities at 0.5, 1.0 and 1.5 M NaCl for each position were similar - or even identical - to those obtained at 0.1M NaCl then no significant deviation from linearity would be observed in the correlation.

The PhAST datasets at four NaCl concentrations were highly correlated and aligned according to x=y (CC = 0.99, Figure 1). This clear result showed that over the considered range, the salt concentration had a marginal effect on the quantum yield of YpY dimer formation and on the average local DNA structure.

It is important to notice that two specific YpY steps are extremely sensitive to laser photo-radiation regardless of the ionic strength (Figure 1, intensity (*I*) > 0.03). These “hotspots” correspond to TpT dinucleotides immediately 5’ of TpA in the two T<u>TT</u>AA segments present in the 601 sequence. According to an NMR study [42], the free TTTAA oligomer is associated with low twists and positive rolls that both favour YpY dimer formation [28] In addition, the two adenines facing the photo-reactive TpT step show uncommon behaviour that includes the resonance broadenings of their H2 and H8 protons as well as an exceptional sensitivity of their ^31^P chemical shifts to temperature changes. The enhanced photo-reactivities associated with these steps could be a further example of the ability of PhAST to detect unusual structural features in DNA.

**Figure 1:**
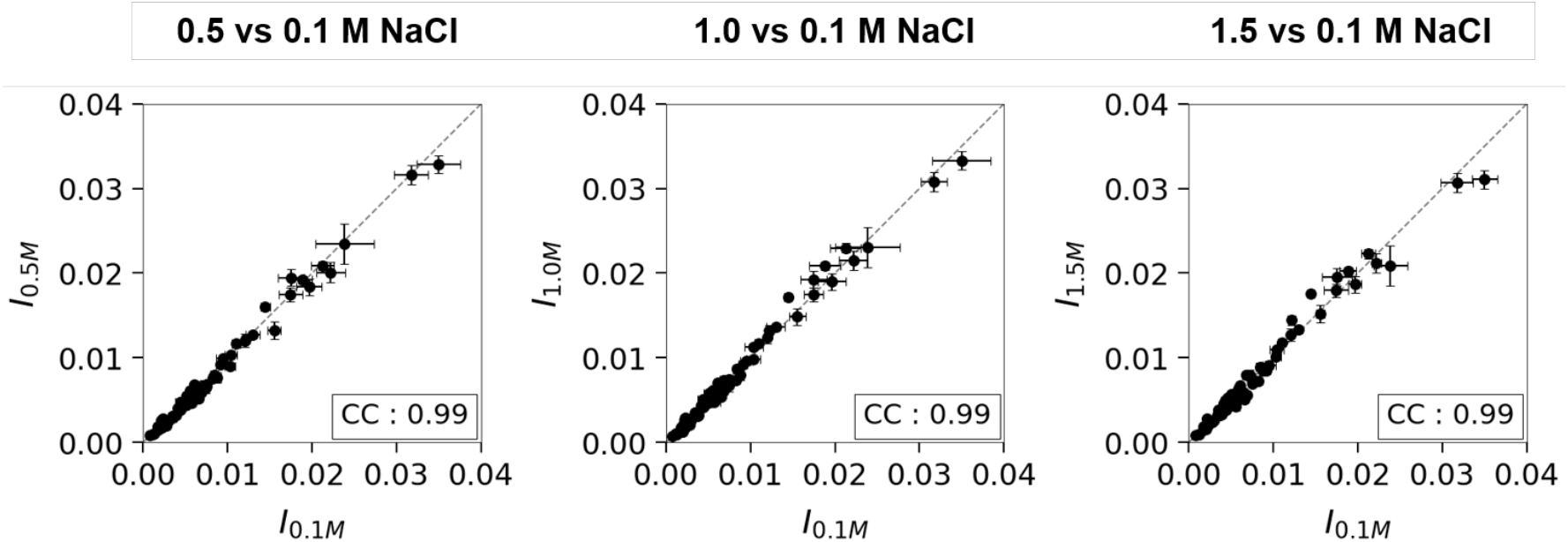
Comparison of PhAST signals at various ionic strengths. The intensity (I) is expressed in terms of normalised peak height associated with each YpY step of the free 601 sequence. The data collected at 0.5, 1.0 and 1.5 M NaCl were systematically compared to those at 0.1 M NaCl. Each point represents the value of I, averaged from 6 experiments related to assembly and disassembly studies; the vertical and horizontal bars are standard errors. The correlation coefficients (CC) are given in boxes in each panel.

In the context of the present work, differences between PhAST signals collected at various ionic strengths can be confidently interpreted in terms of modifications of DNA/histone interactions. The next requirement is to have a precise description of the DNA/histone interface in solution.

### DNA/histone interface

A 1.2 μs explicit solvent molecular dynamics trajectory was recently obtained on a nucleosome containing the 601 sequence and *Xenopus laevis* histones, including large portions of the histone tails (see Materials and Methods). The snapshots, analysed with the VLDM program (see Materials and Methods), provided precise knowledge of DNA/histone contacts in solution [33]. Monitoring of the DNA-protein interface allowed the identification of those DNA regions in which nucleotides of one or the other strand of the double helix interact with the histones H3, H4, H2A or H2B (Figure 2-A). The contribution of each base pair to the interface was also expressed in terms of contact areas involving the DNA and both structured and unstructured histone domains (Figure 2-B). Overall, the contact area associated with (H3-H4)_2_ tetramer represents 54% of the total contact area, thus slightly larger than for H2A-H2B heterodimers. The DNA/histone interactions are remarkably symmetric with respect to the pseudo dyad axis (Figure 2-B) in perfect resonance with the strict symmetry of the histone structured domains; this implies that the DNA sequence, which is not palindromic, has a marginal effect on the interface once the DNA is fully wrapped around the histone and a complete nucleosome is formed.

**Figure 2:**
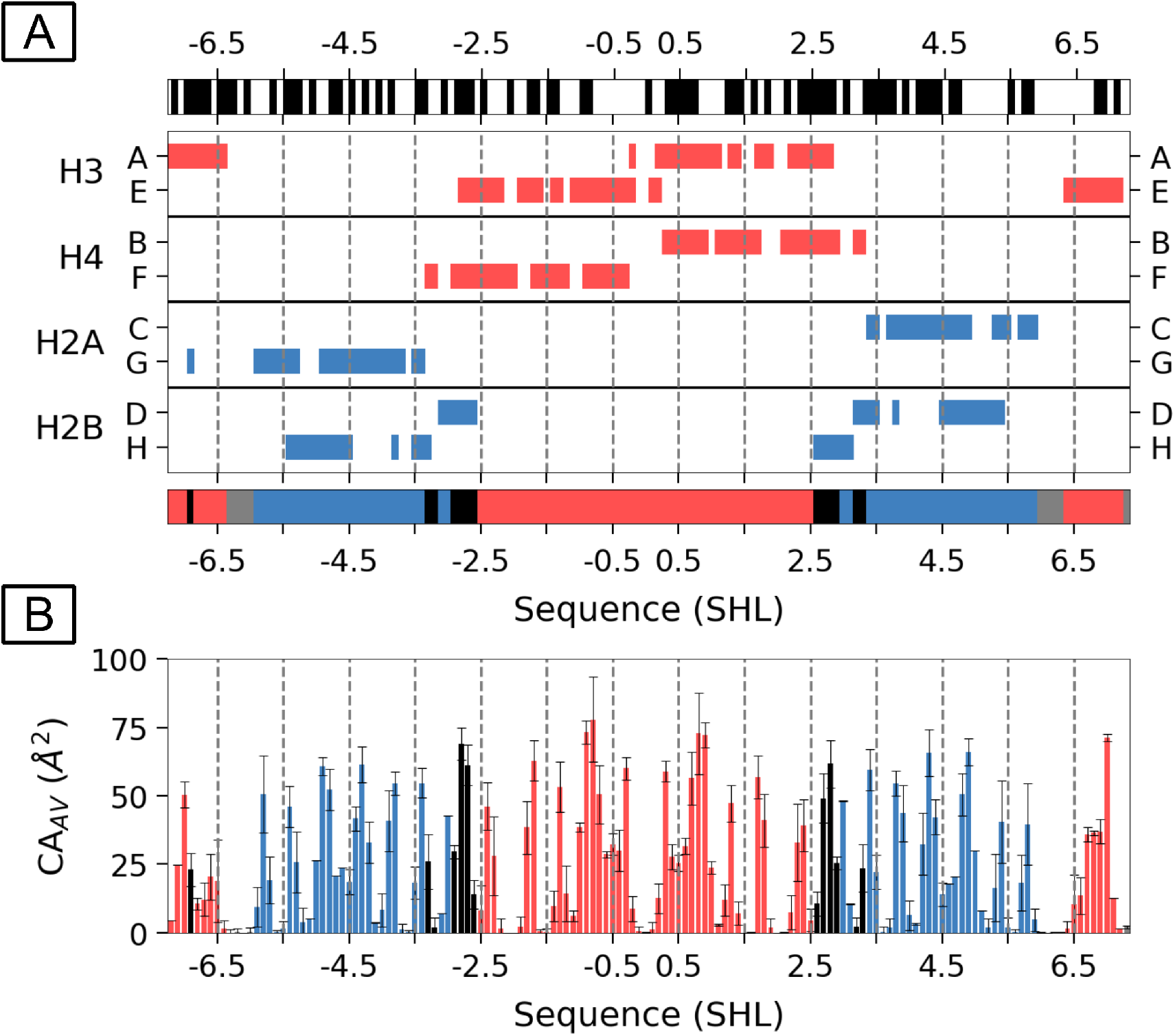
The DNA/histone interface. In both A and B panels, the DNA/histone interface data were extracted and analysed from NCP simulations [33]. **A:** Schematic representation of DNA regions involved in the DNA/histone interface. The nucleotides interacting with the histone structured and unstructured domains are positioned along the 601 sequence, specifying the contacts with the different chains (A, B, C, etc…) of each histone type (H3 and H4 in red, H2A and H2B in blue). The lower banner summarises those regions contacted by H3-H4, H2A-H2B, both H3-H4 and H2A-H2B (black) or none of them (grey). The DNA sequence is labeled by SHL (Super Helical Location, defined in Material and Methods). The dotted lines correspond to intervals of 1 SHL. In the upper banner, the location of YY steps along the two strands of the 601 sequence is pictured by black bars. **B:** Quantification of the DNA/histone interactions. The average contact areas (CA_av_) quantified the DNA/histone interactions at the base pair level of DNA regions, and uses the same color code as in panel A. The vertical error bars are standard deviations over the simulation trajectory. The data presented in panels A and B were extracted from the VLDM analysis of structures from a 1.2 μs simulation of a 601-nucleosome.

On the one hand, there are no DNA regions that escape histone contact apart from a short fragment of 4 bp around SHL ±6.25. However, tracts of Y_n_ (n from 2 to 7) that can form dimers under photo-irradiation are regularly distributed all along the DNA sequence (Figure 2-A). This situation is clearly propitious to the detection of DNA/histone interface modifications using PhAST. Furthermore, differentiating between the effects induced by the binding of the (H3-H4)_2_ tetramer or the H2A-H2B dimers is facilitated by the fact that DNA regions where interactions with H3-H4 and H2A-H2B overlap are extremely limited (Figure 2).

### Assembly and disassembly processes

PhAST was applied to free DNA and nucleosomes at four ionic strengths according to Schema 1. The PhAST signals from assembly experiments were obtained previously [28] and reused here to be compared to those of the disassembly experiments. For comparative purposes they are represented in Figure 3. Changes between PhAST signals of bound and free DNA are quantified by attributing to each YpY position a quantity, log_2_ of Intensity Ratios, defined as log_2_(*IR*) = log_2_ (normalised intensity of a given peak in bound DNA / normalised intensity of the same peak in free DNA), see also Materials and Methods [28] [30]. It should therefore be borne in mind that when the log_2_(*IR*) value is negative this means that the photoreactive signal of the bound DNA ((*I*)_bound DNA_) was less than that for the free DNA ((*I*)_free DNA_).

During both assembly and disassembly experiments, all the YpY steps (Figure 2-A) are associated with PhAST signals. The differences in the log_2_(*IR*) profiles (Figure 3) advocate for the presence of nucleosome intermediate states that are specific of each of the four salt concentrations used. The comparison between log_2_(*IR*) collected at identical ionic strengths during assembly and disassembly experiments is shown in Figure 4.

**Figure 3.**
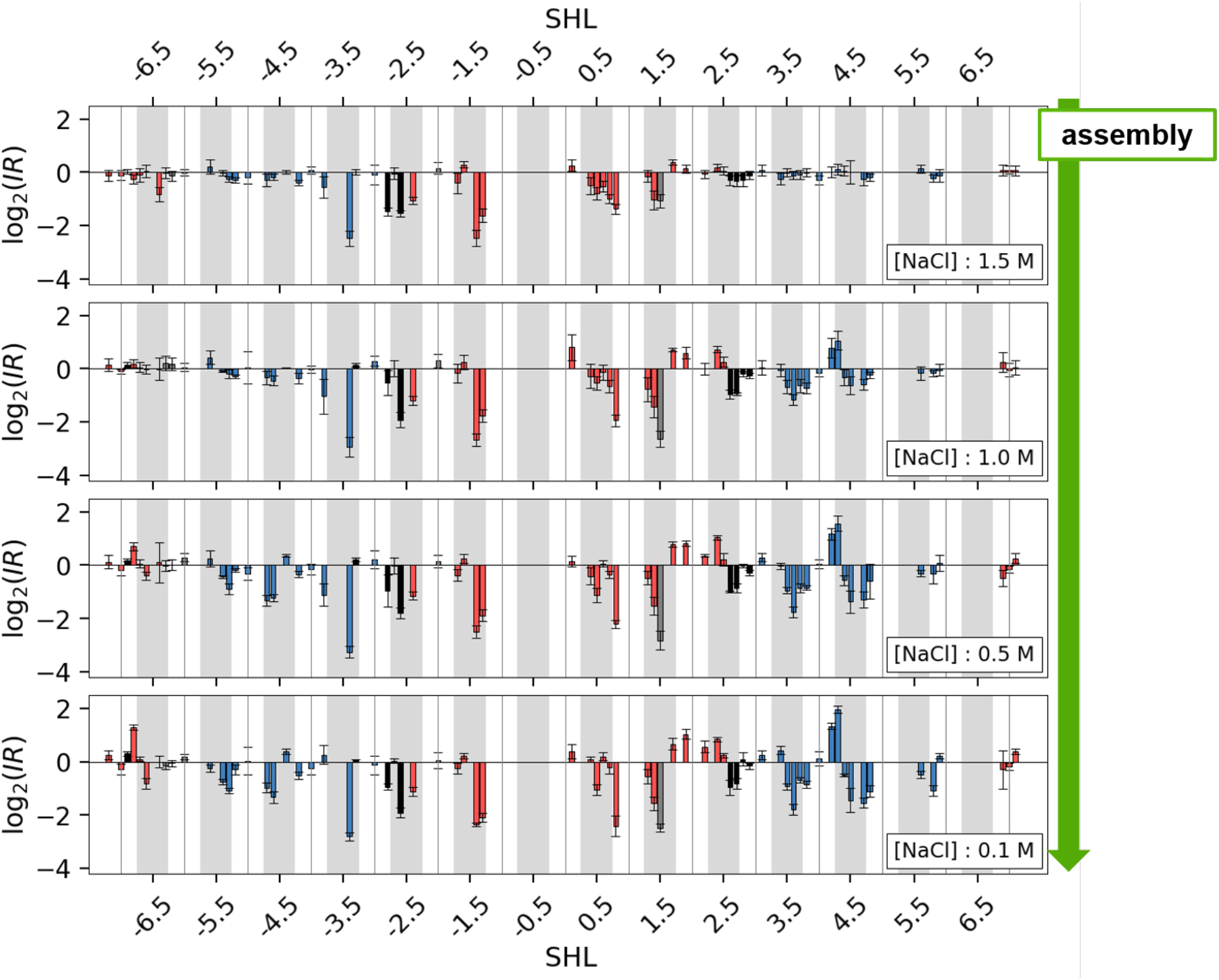
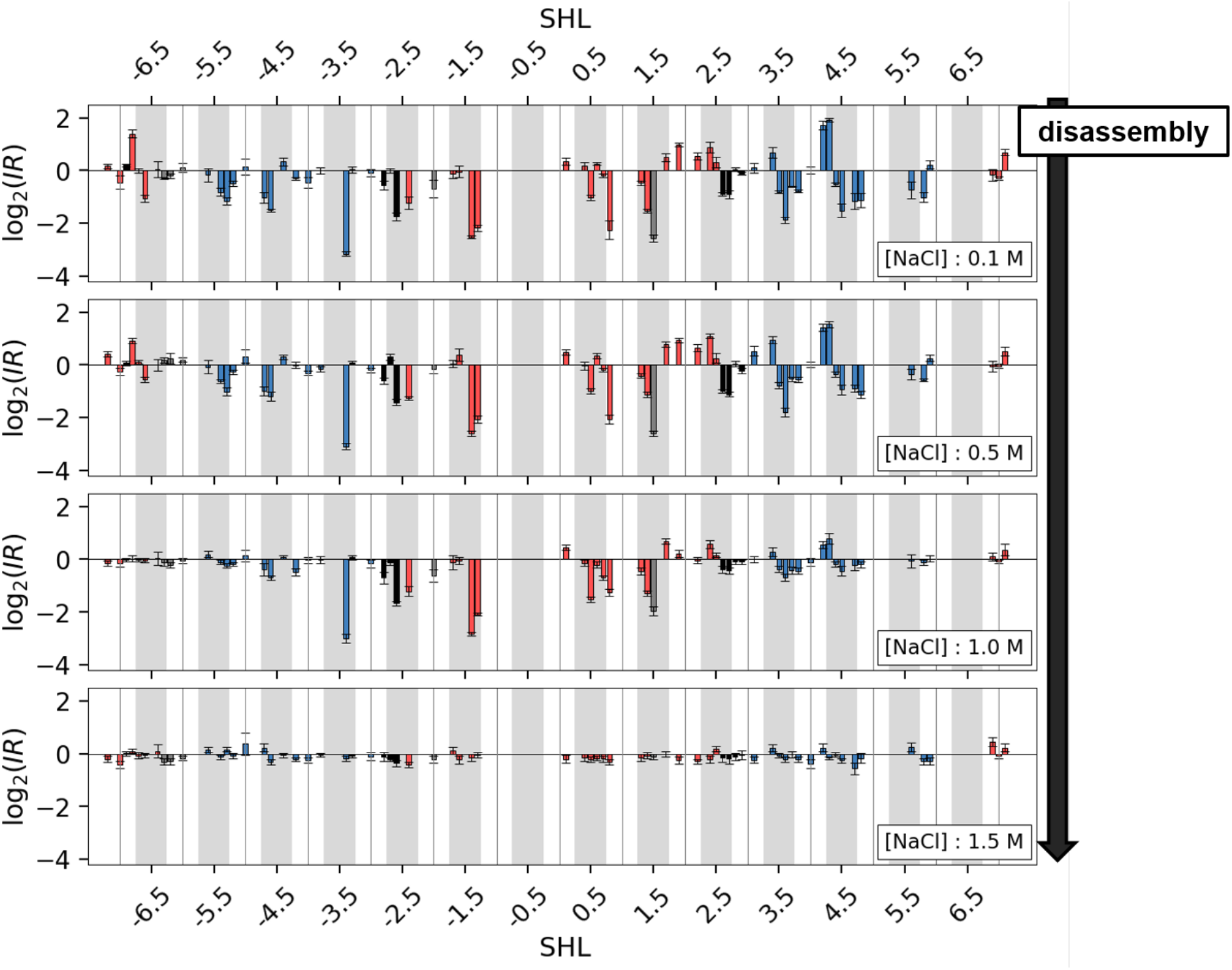
Changes in the probability of YpY dimer formation in DNA during nucleosome assembly and disassembly experiments. PhAST signals are presented in terms of log_2_ of the intensity ratios (IR) along the 601 sequence expressed in SHLs; they are given for decreasing (top panel) or increasing (bottom panel) ionic strengths, as indicated by the green and black arrow respectively. The IR quantities are the ratios calculated between the normalised peak heights of nucleosomal and free DNA at each YpY position (see the text and Materials and Methods); they represent changes in the probability of YpY dimer formation. Red and blue bars correspond to DNA residues involved in the interface with H3-H4 and H2A-H2B, respectively. The black bars correspond to dinucleotides contacted by both H3-H4 and H2A-H2B. Minor-groove inward facing regions observed in the nucleosome structures are represented by grey boxes; they approximatively correspond to the SHL centres. Error bars are standard errors (n≥3, see Material and Methods for details).

**Figure 4:**
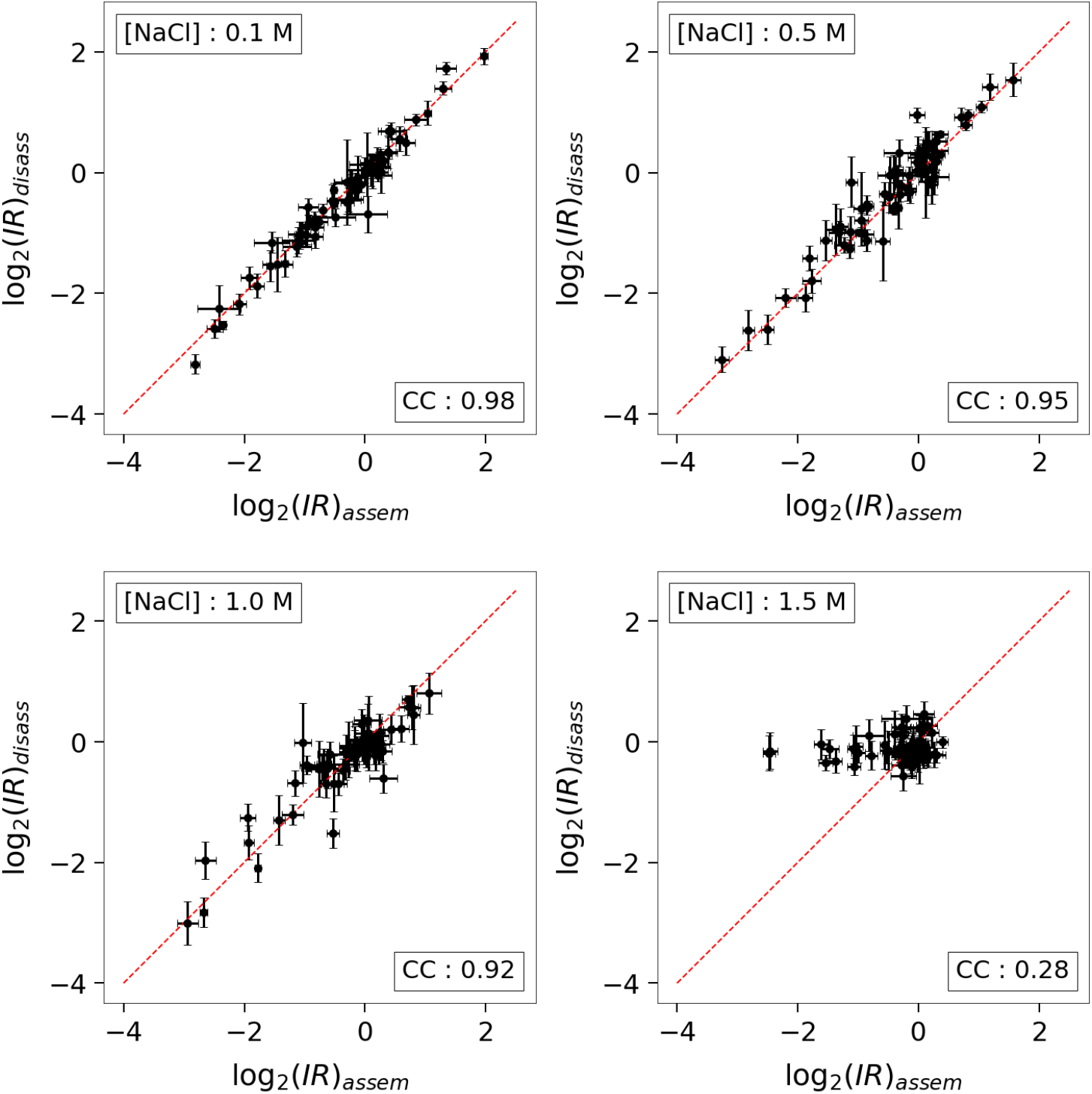
Comparison of PhAST signal changes from assembly and disassembly experiments. Comparison of the log_2_(IR) collected from the assembly (log_2_(IR)_assem_) and disassembly (log_2_(IR)_disass_) experiments at each used ionic strength, i.e. 0.1, 0.5, 1.0 and 1.5 M NaCl. The NaCl concentration and the correlation coefficients (CC) are given in boxes in each panel. log_2_(IR) values for disassembly (ordinate) and assembly (abscissa) are correlated for corresponding positions along the sequence. The red lines represent x=y. Error bars are standard errors (n ≥3, see Material and Methods for details).

At 0.1 M NaCl, the nucleosome was completely formed and stable [28]. Most YpY steps distributed along the whole DNA length showed significant values of log_2_(*IR*) (Figure 3 and Figure 4, top left). The PhAST signal profile at 0.1 M NaCl represents a signature of the structural effect of the histone octamer on the DNA in a fully formed nucleosome. As previously shown [28], a decrease in the probability of formation of a given YpY dimer ((*I*)_bound DNA_ < (*I*)_free DNA_, log_2_(*IR*) < 0) is produced when, from free to bound, the value of roll changes i) from a positive value to a less positive or negative value or ii) from a negative value to a more negative value. The interpretation of an increase in probability of YpY dimer formation arises from the same scheme, substituting negative by positive and positive by negative. So, the sign of log_2_(*IR*) finally relates to the structural nature of histone-induced changes.

The similarity between assembly and disassembly experiments at 0.1 M NaCl globally persisted at 0.5 M but began to break down at 1.0 M and finally disappeared at 1.5 M (Figure 4). To further characterise the events occurring during assembly and disassembly experiments at ionic strengths other than 0.1 M, we calculated R_log2(*IR*)_, defined as the ratio between |log_2_(*IR*)| at a given ionic strength and ļlog_2_(*IR*)ļ at 0.1 M, the PhAST signal associated with an intact nucleosome, which we therefore use as reference. Note that the R_log2(*IR*)_ quantity does not take into account the positive or negative sign of PhAST signals that only have a structural significance as mentioned previously. Low R_log2(*IR*)_ values relate to weak DNA/histone interactions with the extreme, and ideal case of R_log2(*IR*)_ = 0 for a DNA totally free of histones; high R_log2(*IR*)_ values reflect robust DNA/histone contacts, a DNA fully wrapped around the histone octamer ideally corresponding to R_log2(*IR*)_ = 1. R_log2(*IR*)_ values were calculated by summing |log_2_(*IR*)| values over 10-bp fragments centred on SHLs ±0.5, ±1.5, etc… according to the DNA/histone interaction pattern (Figure 2). The result of this analysis is summarized in Figure 5 and illustrated by cartoons of molecular models in Figures S2-1 and S2-2.

**Figure 5:**
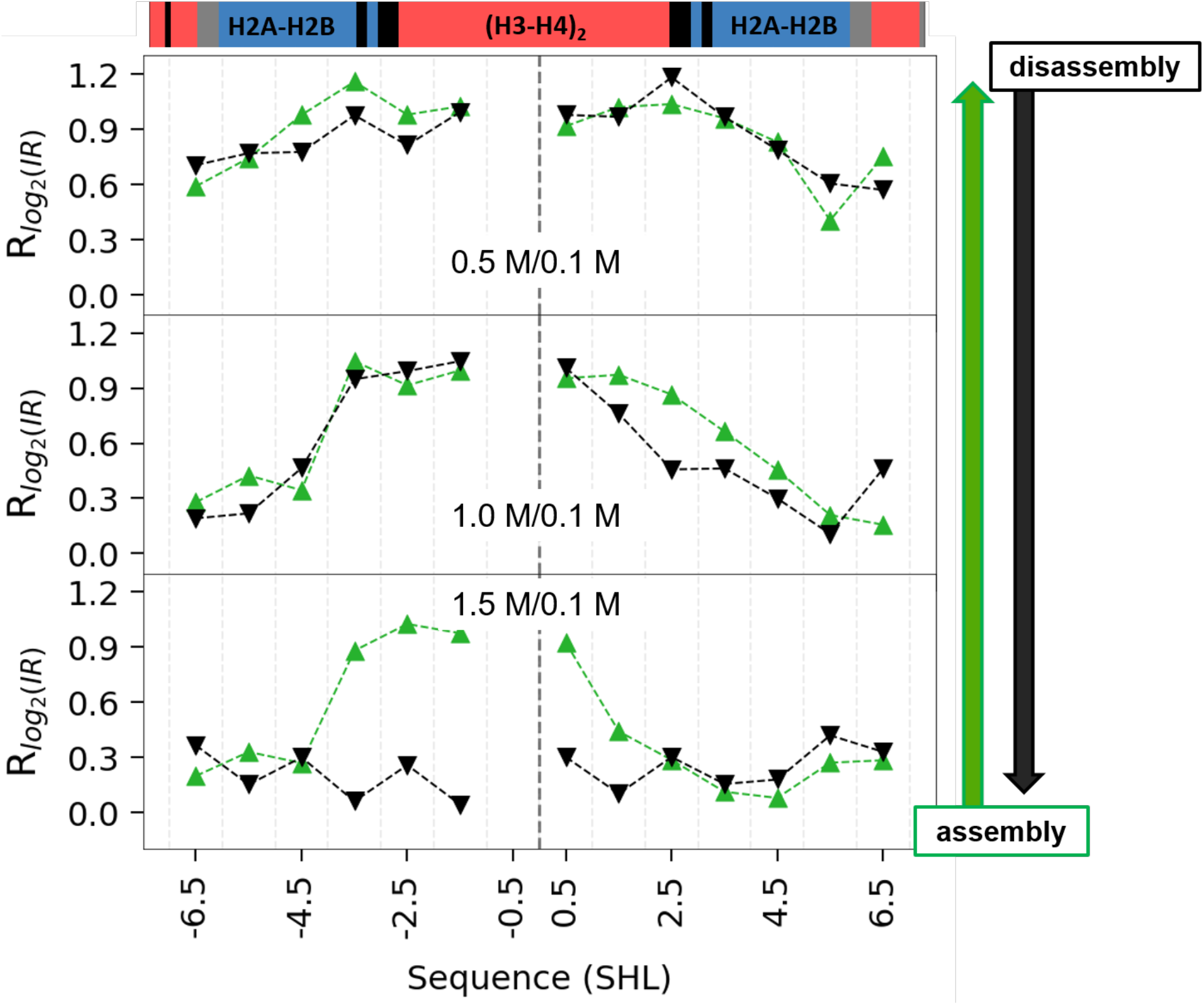
Changes in PhAST signals by SHLs, in assembly and disassembly experiments. The changes of PhAST signals between 0.1 M and the other salt concentration are expressed by the ratios (R_log2(IR)_) between the absolute values of log_2_(IR) at a given ionic strength and log_2_(IR) at 0.1 M, calculated by SHL. High and low R_log2(IR)_ values indicate increasing and decreasing effects of DNA/histone interactions on DNA shape. The data were sorted and averaged by SHL. The assembly (green triangles) and disassembly (black inverted-triangles) data are compared at each ionic strength. The vertical dashed line indicates SHL=0. The boxes at the top specify those histones that interact: H3 and H4 in red, H2A and H2B in blue, both H3-H4 and H2A-H2B in black and where none are involved in grey.

In both assembly and disassembly experiments, the resemblances between PhAST signals at 0.5 M and 0.1 M (Figure 5) were compatible with the dominant presence of stable DNA/(H3-H4)_2_ interactions around the DNA pseudodyad. The tendency of R_log2(*IR*)_ values to decrease at the DNA extremities at 0.5 M (Figure 5) indicated that H2A-H2B and especially H2A(C)-H2B(D) around SHL +5.5 were more loosely bound to the DNA than (H3-H4)_2_. This instability also concerns the H3 tails that contact the DNA at SHLs ± 6.5.

The comparison of PhAST signals at 1 M and 0.1 M revealed a more complex situation. The low R_log2(*IR*)_ values associated with the two peripheral DNA regions (Figure 5) signalled a severe weakening of the interface between DNA and the two (H2A-H2B) dimers in both assembly and disassembly experiments. In the DNA centre, the two sides around the pseudodyad responded differently to photo-irradiation, generating a clear asymmetry (Figure 5). On the 3’ side, PhAST signal changes expanded from the 3’ end to near the centre (Figure 5), reflecting alterations in interactions with H3 and H4, enhanced in the disassembly experiments (illustrated by the cartoons in Figure 6-A). On the 5’ side, the high R_log2(*IR*)_ values observed from SHL 0 to SHL −3.5 (Figure 5) implied that the ~35 bp region 5’ of the pseudodyad remained tightly bound to H3 and H4 in both association and dissociation experiments (Figure S2). Within this same region, the abrupt drop of R_log2(*IR*)_ values between SHL −3.5 and SHL −4.5 (Figure 5) also indicated that the integrity of the H3-H4 contacts was not affected by the serious deterioration of DNA/H2A-H2B dimer interactions present at SHLs −3.5 and −2.5, where the four histones meet (Figure 2).

According to PhAST, assembly and disassembly intermediate states at 1.5 M only, shared the critical deterioration (Figures 4 and 5), or even the absence, of interactions of DNA with the two H2A-H2B dimers (Figure 5). 1.5 M corresponds to the first step of nucleosome assembly. In this case, there is again a strong asymmetry with respect to the DNA centre (Figures 4, 5 and S2) that matches with an octamer (H3-H4)_2_ still firmly attached to half of its DNA site (Figure 6-B). In contrast, at the same salt concentration, the last step of the disassembly process is described by log_2_(*IR*) values approaching zero (Figure 3) and low R_log2(*IR*)_ values (Figure 5) all along the DNA. Clearly here, the asymmetry is totally lost. The close resemblance with what happens with the free DNA is consistent with a massive histone release (Figures 5 and 6-B), in agreement with experiments using high precision FRET approaches [2].

**Figure 6:**
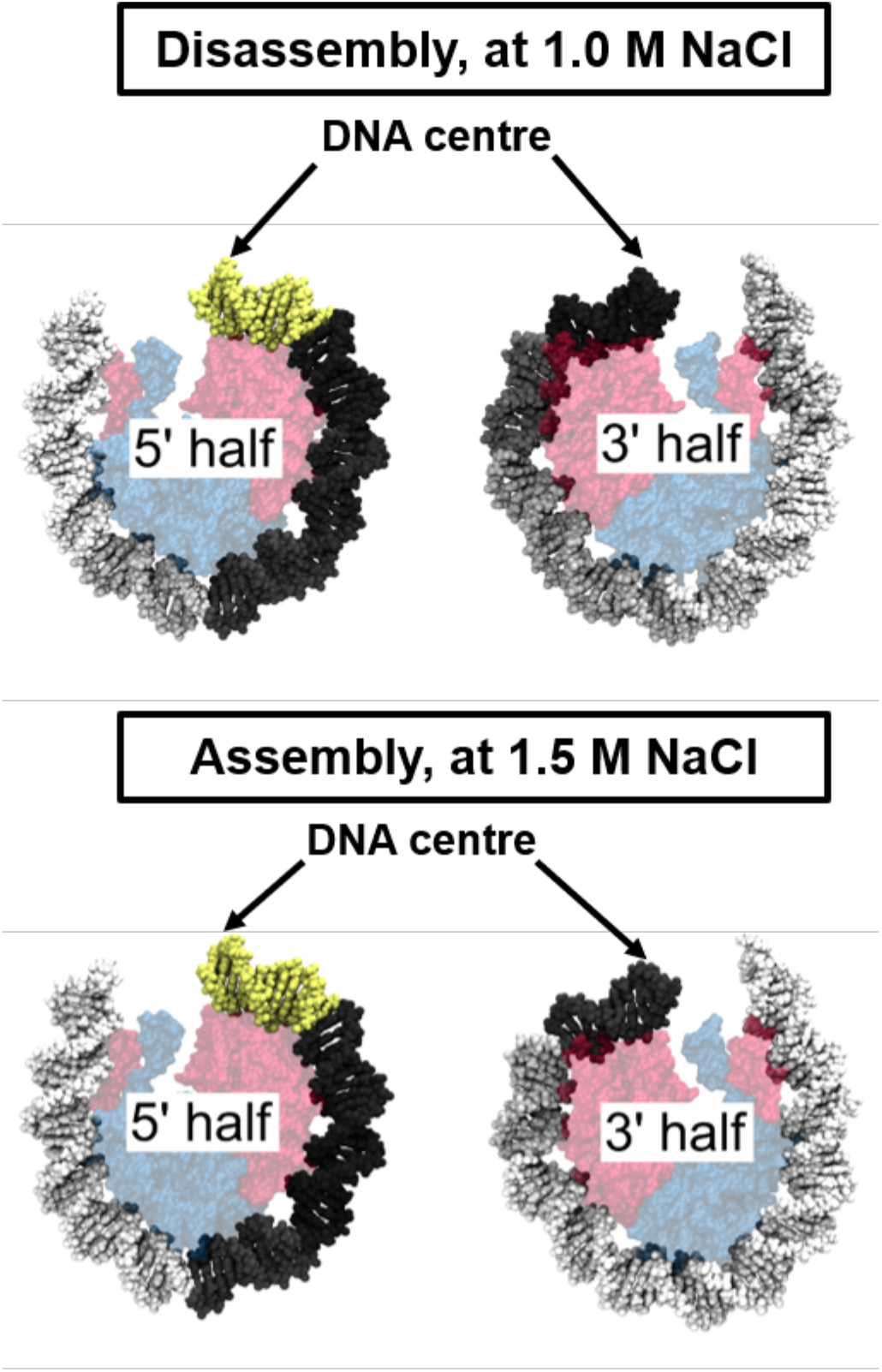
Representation of asymmetries in the DNA/histone interactions on nucleosome models according to PhAST experiments. In the panels A and B, the DNA double helix was coloured with a gradient of grey, from light to dark grey for low and high R_log2(IR)_ values (see Figure 5). Low and high R_log2(IR)_ values were interpreted as weak and strong DNA/histone interactions. The double helix in yellow corresponds to the DNA region devoid of YpY steps. The nucleosome DNA was split into two parts containing either the 5’ or 3’ half of the 601 sequence. Histones are represented by semi-transparent models (H3-H4 in red and H2A-H2B in blue) to reflect their location in the intact nucleosome. For clarity, the histone tails are not represented. Comparison of the PhAST derived data R_log2(IR)_ along the 5’ (left) and 3’ (right) DNA halves **A**: at 1.0 M NaCl in the experiments of nucleosome dissociation and **B**: at 1.5 M NaCl in the experiments of nucleosome association.

## Discussion

The main aim of this study was to provide a comparison of nucleosome formation and dissociation pathways under the same conditions. We used the PhAST approach that is non-invasive and does not require the use of chemical modifications. The unambiguous detection of nucleosome intermediate states in terms of base pair location along the whole nucleosomal DNA is one of the major advantages of PhAST over existing techniques used to investigate transitions in nucleosome organisation.

Clearly, multiple intermediate states are formed sequentially during both nucleosome assembly and disassembly. We infer that during assembly, binding takes place from the DNA centre and progresses towards the extremities whereas during disassembly, dissociation develops in the direction from the extremities to the centre. These directionally inverse progressions of histone interactions are not strictly symmetrical with respect to the pseudo dyad axis, an observation that merits a more detailed discussion, in particular regarding the DNA sequence effects underlying the changes of the DNA/histone interactions detected by PhAST.

At 0.5 M NaCl, the PhAST signals at the two DNA extremities matched with DNA breathing, probably enhanced at the 3’ extremity as previously suggested [22]. When the ionic strength exceeded 0.5 M, the DNA/H2A-H2B interfaces were increasingly weakened so that differences between 5’ and 3’ extremities became undetectable. As previously explained [28] [33], electrostatic interactions are expected to be weakened at high ionic strength and consequently hydrophobic contacts should play a more important role at the interfaces. By the same logic, the DNA/H2A-H2B interfaces are disadvantaged because of their poor hydrophobic character and the relatively small area of contact compared to those of the DNA/H3-H4 interface [33].

A novel aspect of our work involves the 70-bp region around the pseudo dyad, more precisely those segments covering SHLs −3.5 → 0.5 (in the 5’ half) and SHLs 0.5 → 3.5 (in the 3’ half). PhAST revealed their asymmetric behaviour at 1.5 M NaCl for assembly and at 1 M for disassembly (Figure 6): in both cases the 5’ side was clearly more favourable to H3-H4 anchoring than the 3’ side.

Concerning assembly, previous results from our TRX annotation provided a quantitative characterisation of the intrinsic structural variability in B-DNA, at the dinucleotide level [24]. In terms of what is predicted from the TRX analysis on free DNA, the 601 sequence would consist of asymmetric halves [24] [42]. From the 5’ extremity to SHL 2, the alternation of stiff and flexible 5-bp segments was found to perfectly coincide with the periodic, sinusoidal variations of the structural descriptors of the DNA wrapped in a nucleosome. This correspondence was strongly reduced or even disappeared in regions from SHL 2.5 to the 3’ extremities, as in other sequences of lower affinity for the histones. It was concluded from these studies that the ability of the 601 sequence to form nucleosomes originates from the intrinsic structural and dynamic properties of the whole 5’ half extended to the segment just 3’ of the dyad (SHLs −7 to 2.5), which limit the cost of DNA wrapping. This idea also provided a convincing explanation for the asymmetry that we observed here during assembly using PhAST [28].

This 5’ vs 3’ side asymmetry involving the central 70 bp is also present during disassembly. A first hypothesis is that this asymmetry signals variations of the DNA-histone interface strength along the DNA in the intact, complete nucleosome, as occurs with H2A-H2B. However, this is clearly not the case since analysis of a stable nucleosome in solution showed that the concerned segments, symmetrically located with respect to the pseudo dyad, interact equally with (H3-H4)_2_ (Figure 2) [33]. We suggest that the disassembly pathway exploits the most stressed regions in a manner analogous to the relaxation of a stiff spring upon release of constraints. Indeed, our work indicates that those DNA regions that are particularly refractive to nucleosome formation are also the first to break their interactions with the histones.

In summary, the PhAST approach provides a fine picture of the propensities of DNA and histones to interact that, in turn, allows the decryption of the main nucleosome intermediate states present during assembly and disassembly. Transitions between the relatively simple ensembles of octasomes, hexasomes or tetrasomes described hitherto in the literature are in fact more subtle than expected; we obtain snapshots of a continuous process of gradual rearrangements leading to the final structure. Although under the conditions used here, assembly and disassembly pathways do not correspond to strictly invertible schemes, both are extremely sensitive to the DNA sequence. Strikingly, our detailed analysis provides unambiguous evidence for the role of the DNA sequence in determining the relative stability of these intermediates, through its effect on local DNA structure and stiffness. We suggest that DNA sequence contributes to nucleosome positioning mainly through differential stabilisation of these intermediates, rather than of fully formed particles. The ability to reliably investigate these processes makes PhAST a promising tool for the study of how DNA sequence affects chromatin remodelling *in vivo*, and creates exciting opportunities to design DNA sequences with varying affinities for histones that could be useful to fully explore the complex relationship between DNA sequence and nucleosome positioning. Currently, even though it is of course difficult to extrapolate these results to the *in vivo* situation, it seems reasonable to expect that such DNA sequence effects modulate chromatin remodelling in conjunction with numerous trans factors such as ATP-dependent proteins that regulate binding, release and sliding of nucleosomes.

## Acknowledgements

Andrew Travers is warmly thanked for having initiated and guided this study and whose ideas are a continuous thread throughout. The authors also thank Christophe Oguey for helpful advice. This manuscript is dedicated to the memory of Jorge Langowski whose animated and instructive discussions with MB strongly challenged and influenced the ideas expressed in this manuscript.

## Authors contributions

C.N. and M.B. conceptualized the study. A.H. performed PhAST experiments. R.R. and M.P. performed the analyses. B.H. and M.B. collected the bibliography and wrote the paper. All authors discussed the results and commented on the paper.

## References

1 Park, Y. J., Dyer, P. N., Tremethick, D. J. and Luger, K. (2004) A new fluorescence resonance energy transfer approach demonstrates that the histone variant H2AZ stabilizes the histone octamer within the nucleosome. J Biol Chem. 279, 24274–24282

2 Gansen, A., Felekyan, S., Kuhnemuth, R., Lehmann, K., Toth, K., Seidel, C. A. M. and Langowski, J. (2018) High precision FRET studies reveal reversible transitions in nucleosomes between microseconds and minutes. Nat Commun. 9, 4628

3 Hall, M. A., Shundrovsky, A., Bai, L., Fulbright, R. M., Lis, J. T. and Wang, M. D. (2009) High-resolution dynamic mapping of histone-DNA interactions in a nucleosome. Nat Struct Mol Biol. 16, 124–129

4 Thuy T.M. Ngo, Q. Z., Ruobo Zhou, Jaya G. Yodh, and Taekjip Ha. (2015) Asymmetric Unwrapping of Nucleosomes under Tension Directed by DNA Local Flexibility. Cell. 160, 1135–1144

5 Sheinin, M. Y., Li, M., Soltani, M., Luger, K. and Wang, M. D. (2013) Torque modulates nucleosome stability and facilitates H2A/H2B dimer loss. Nat Commun. 4, 2579

6 Tomschik, M., Zheng, H., van Holde, K., Zlatanova, J. and Leuba, S. H. (2005) Fast, long-range, reversible conformational fluctuations in nucleosomes revealed by single-pair fluorescence resonance energy transfer. Proc Natl Acad Sci U S A. 102, 3278–3283

7 Koopmans, W. J., Buning, R., Schmidt, T. and van Noort, J. (2009) spFRET using alternating excitation and FCS reveals progressive DNA unwrapping in nucleosomes. Biophys J. 97, 195–204

8 Miyagi, A., Ando, T. and Lyubchenko, Y. L. (2011) Dynamics of nucleosomes assessed with time-lapse high-speed atomic force microscopy. Biochemistry. 50, 7901–7908

9 Lyubchenko, Y. L. (2014) Nanoscale Nucleosome Dynamics Assessed with Time-lapse AFM. Biophys Rev. 6, 181–190

10 Bilokapic, S., Strauss, M. and Halic, M. (2018) Histone octamer rearranges to adapt to DNA unwrapping. Nat Struct Mol Biol. 25, 101–108

11 Bohm, V., Hieb, A. R., Andrews, A. J., Gansen, A., Rocker, A., Toth, K., Luger, K. and Langowski, J. (2011) Nucleosome accessibility governed by the dimer/tetramer interface. Nucleic Acids Res. 39, 3093–3102

12 Hoch, D., A; J,J; Stratton, and Gloss, L,M. (2007) Protein-Protein Förster Resonance Energy Transfer Analysis of Nucleosome Core Particles Containing H2A and H2A.Z. JMB. 371, 971–988

13 Gansen, A., Valeri, A., Hauger, F., Felekyan, S., Kalinin, S., Toth, K., Langowski, J. and Seidel, C. A. (2009) Nucleosome disassembly intermediates characterized by single-molecule FRET. Proc Natl Acad Sci U S A. 106, 15308–15313

14 Toth, K. B., V; Sellmann, C; Danner, M; Hanne, J; Berg, M; and Barz, I. G., A; Langowski, J. (2013) Histone-and DNA Sequence-DependentStability of Nucleosomes Studied by Single-Pair FRET. Cytometry Part A. 83A, 839–846

15 Lee, J. and Lee, T. H. (2017) Single-Molecule Investigations on Histone H2A-H2B Dynamics in the Nucleosome. Biochemistry. 56, 977–985

16 Chen, Y., Tokuda, J. M., Topping, T., Sutton, J. L., Meisburger, S. P., Pabit, S. A., Gloss, L. M. and Pollack, L. (2014) Revealing transient structures of nucleosomes as DNA unwinds. Nucleic Acids Res. 42, 8767–8776

17 Chen, Y., Tokuda, J. M., Topping, T., Meisburger, S. P., Pabit, S. A., Gloss, L. M. and Pollack, L. (2017) Asymmetric unwrapping of nucleosomal DNA propagates asymmetric opening and dissociation of the histone core. Proc Natl Acad Sci U S A. 114, 334–339

18 Kelbauskas, L. C., N; Bash, R; Yidh, J; Woodbury, N. Lohr,D. (2007) Sequence-Dependent Nucleosome Structure and Stability Variations Detected by Förster Resonance Energy Transfer†. Biochemistry. 46, 2239–2248

19 Arimura, Y., Tachiwana, H., Oda, T., Sato, M. and Kurumizaka, H. (2012) Structural analysis of the hexasome, lacking one histone H2A/H2B dimer from the conventional nucleosome. Biochemistry. 51, 3302–3309

20 Kato, D. O., A. Arimura, Y. Mizukami,Y., Horikoshi, N. S., K. Akashi, S. Nishimura,Y., Park, S.-Y. N., J. Maehara, K. Ohkawa, Y. and Matsumoto, A. K., H. Inoue, R. Sugiyama, H, Kurumizaka, H. (2017) Crystal structure of the overlapping dinucleosome composed of hexasome and octasome. Science. 356, 205–208

21 Mauney, A. W., Tokuda, J. M., Gloss, L. M., Gonzalez, O. and Pollack, L. (2018) Local DNA Sequence Controls Asymmetry of DNA Unwrapping from Nucleosome Core Particles. Biophys J. 115, 773–781

22 Chua, E. Y., Vasudevan, D., Davey, G. E., Wu, B. and Davey, C. A. (2012) The mechanics behind DNA sequence-dependent properties of the nucleosome. Nucleic Acids Res. 40, 6338–6352

23 Luger, K., Mader, A. W., Richmond, R. K., Sargent, D. F. and Richmond, T. J. (1997) Crystal structure of the nucleosome core particle at 2.8 A resolution. Nature. 389, 251–260

24 Heddi, B., Oguey, C., Lavelle, C., Foloppe, N. and Hartmann, B. (2010) Intrinsic flexibility of B-DNA: the experimental TRX scale. Nucleic Acids Res. 38, 1034–1047

25 Sullivan, S. A. a. L., D. (2003) Characterization of Sequence Variability in NucleosomeCore Histone Folds. PROTEINS. 52, 454–465

26 Worcel, A., Han, S. and Wong, M. L. (1978) Assembly of newly replicated chromatin. Cell. 15, 969–977

27 Cremisi, C. and Yaniv, M. (1980) Sequential assembly of newly synthesized histones on replicating SV40 DNA. Biochem Biophys Res Commun. 92, 1117–1123

28 Hatakeyama, A., Hartmann, B., Travers, A., Nogues, C. and Buckle, M. (2016) High-resolution biophysical analysis of the dynamics of nucleosome formation. Scientific Reports. 6, 27337

29 Buckle, M., Geiselmann, J., Kolb, A. and Buc, H. (1991) Protein-DNA cross-linking at the lac promoter. Nucleic Acids Research. 19, 833–840

30 Brach, K., Hatakeyama, A., Nogues, C., Olesiak-Banska, J., Buckle, M. and Matczyszyn, K. (2018) Photochemical analysis of structural transitions in DNA liquid crystals reveals differences in spatial structure of DNA molecules organized in liquid crystalline form. Sci Rep. 8, 4528

31 Imeddourene, A. B., Xu, X., Zargarian, L., Oguey, C., Foloppe, N., Mauffret, O. and Hartmann, B. (2016) The intrinsic mechanics of B-DNA in solution characterized by NMR. Nucleic Acids Res

32 Andrews, A. J. and Luger, K. (2011) Nucleosome Structure(s) and Stability: Variations on a Theme. Annu Rev Biophys. 40, 99–117

33 Elbahnsi, A., Retureau, R., Baaden, M., Hartmann, B. and Oguey, C. (2018) Holding the Nucleosome Together: A Quantitative Description of the DNA-Histone Interface in Solution. J Chem Theory Comput. 14, 1045–1058

34 Dumuiskervabon, A., Encontre, I., Etienne, G., Jaureguiadell, J., Mery, J., Mesnier, D. and Parello, J. (1986) A Chromatin Core Particle Obtained by Selective Cleavage of Histones by Clostripain. Embo Journal. 5, 1735–1742

35 Morales, V. and Richard-Foy, H. (2000) Role of histone N-terminal tails and their acetylation in nucleosome dynamics. Molecular and Cellular Biology. 20, 7230–7237

36 Hart, K., Foloppe, N., Baker, C. M., Denning, E. J., Nilsson, L. and MacKerell, A. D. (2012) Optimization of the CHARMM Additive Force Field for DNA: Improved Treatment of the BI/BII Conformational Equilibrium. Journal of Chemical Theory and Computation. 8, 348–362

37 Retureau, R., Oguey, C., Mauffret, O. and Hartmann, B. (2019) Structural Explorations of NCp7-Nucleic Acid Complexes Give Keys to Decipher the Binding Process. J Mol Biol. 431, 1966–1980

38 Schildkraut, C. and Lifson, S. (1965) Dependence of Melting Temperature of DNA on Salt Concentration. Biopolymers. 3, 195–+

39 Borochov, N., Eisenberg, H. and Kam, Z. (1981) Dependence of DNA Conformation on the Concentration of Salt. Biopolymers. 20, 231–235

40 Baumann, C. G. S., S B. Bloomfeld, V A. Bustamante, C. (1997) Ionic effects on the elasticity of single DNA molecules. Proc. Natl. Acad. Sci. USA. 94, 6185–6190

41 Pan, S., At Nguyen, D., Sridhar, T., Sunthar, P. and Ravi Prakash, J. (2014) Universal solvent quality crossover of the zero shear rate viscosity of semidilute DNA solutions. Journal of Rheology. 58, 339–368

42 Xu, X., Ben Imeddourene, A., Zargarian, L., Foloppe, N., Mauffret, O. and Hartmann, B. (2014) NMR studies of DNA support the role of pre-existing minor groove variations in nucleosome indirect readout. Biochemistry. 53, 5601–5612

